# Modulation of taste sensitivity by the olfactory system in *Drosophila*

**DOI:** 10.1101/2021.03.30.437740

**Authors:** Pierre Junca, Molly Stanley, Pierre-Yves Musso, Michael D Gordon

## Abstract

An animal’s sensory percepts are not raw representations of the outside world. Rather, they are constructs influenced by many factors including the species, past experiences, and internal states. One source of perceptual variability that has fascinated researchers for decades is the effect of losing one sensory modality on the performance of another^1^. Typically, dysfunction of one sense has been associated with elevated function of others, creating a type of sensory homeostasis^2^. For example, people with vision loss have been reported to demonstrate enhanced tactile and auditory functions, and deafness has been associated with heightened attention to visual inputs for communication^3,4^. By contrast, smell and taste—the two chemosensory modalities—are so intrinsically linked in their contributions to flavor that loss of smell is often anecdotally reported as leading to deficiencies in taste^5–8^. However, human studies specifically examining taste are mixed and generally do not support this widely-held belief, and data from animal models is largely lacking^9^. Here, we examine the impact of olfactory dysfunction on taste sensitivity in *Drosophila melanogaster*. We find that partial loss of olfactory input (hyposmia) dramatically enhances flies’ sensitivity to both appetitive (sugar, low salt) and aversive (bitter, high salt) tastes. This taste enhancement is starvation-independent and occurs following suppression of either first- or second-order olfactory neurons. Moreover, optogenetically increasing olfactory inputs reduces taste sensitivity. Finally, we observed that taste enhancement is not encoded in the activity of peripheral gustatory sensory neurons, but is associated with elevated sugar responses in protocerebrum anterior medial (PAM) dopaminergic neurons of the mushroom bodies. These results suggest a level of homeostatic control over chemosensation, where flies compensate for lack of olfactory input by increasing the salience of taste information.

## Results and Discussion

Fruit flies rely heavily on chemosensation to find, select, and consume food. Although smell operates to detect food volatiles over long ranges and taste occurs only on contact, there are many scenarios where the two senses are active simultaneously^10,11^. Yet, aside from the impact of coincident olfactory and gustatory inputs on learning, we know little about how one chemical sense influences the other^12,13^. To explore this question, we examined *Drosophila* mutants for *Odorant receptor co-receptor* (*Orco*), which are hyposmic due to complete loss of OR-dependent olfaction. *Orco* mutants lack both spontaneous and evoked firing in OR-expressing olfactory receptor neurons (ORNs), and therefore receive dramatically lower levels of total input to the olfactory system^14,15^. We began by testing the effect of hyposmia on feeding by employing the FlyPAD^16^, which measures preference between two food sources presented in 1% agar (Figure 1A). In this assay, control *w*^1118^ flies showed little to no preference for 1 mM sucrose over water after 24 hours of starvation. However, three different allelic combinations of *Orco* mutants displayed significantly higher preference for this low sucrose concentration than controls (Figures 1B,S1A). This phenotype was replicated upon silencing of Orco neurons with conditional expression of Kir2.1, demonstrating that suppressed activity in the olfactory system leads to heightened sugar feeding (Figures 1C,S1B). To further characterize the nature of this feeding enhancement, we measured sucrose preference across a series of concentrations. This experiment revealed that hyposmic flies begin to exhibit a significant preference for sucrose over water at substantially lower concentrations than controls (Figure 1D,S1C). Thus, loss of olfactory inputs appears to dramatically increase sweet taste sensitivity.

**Figure 1.**
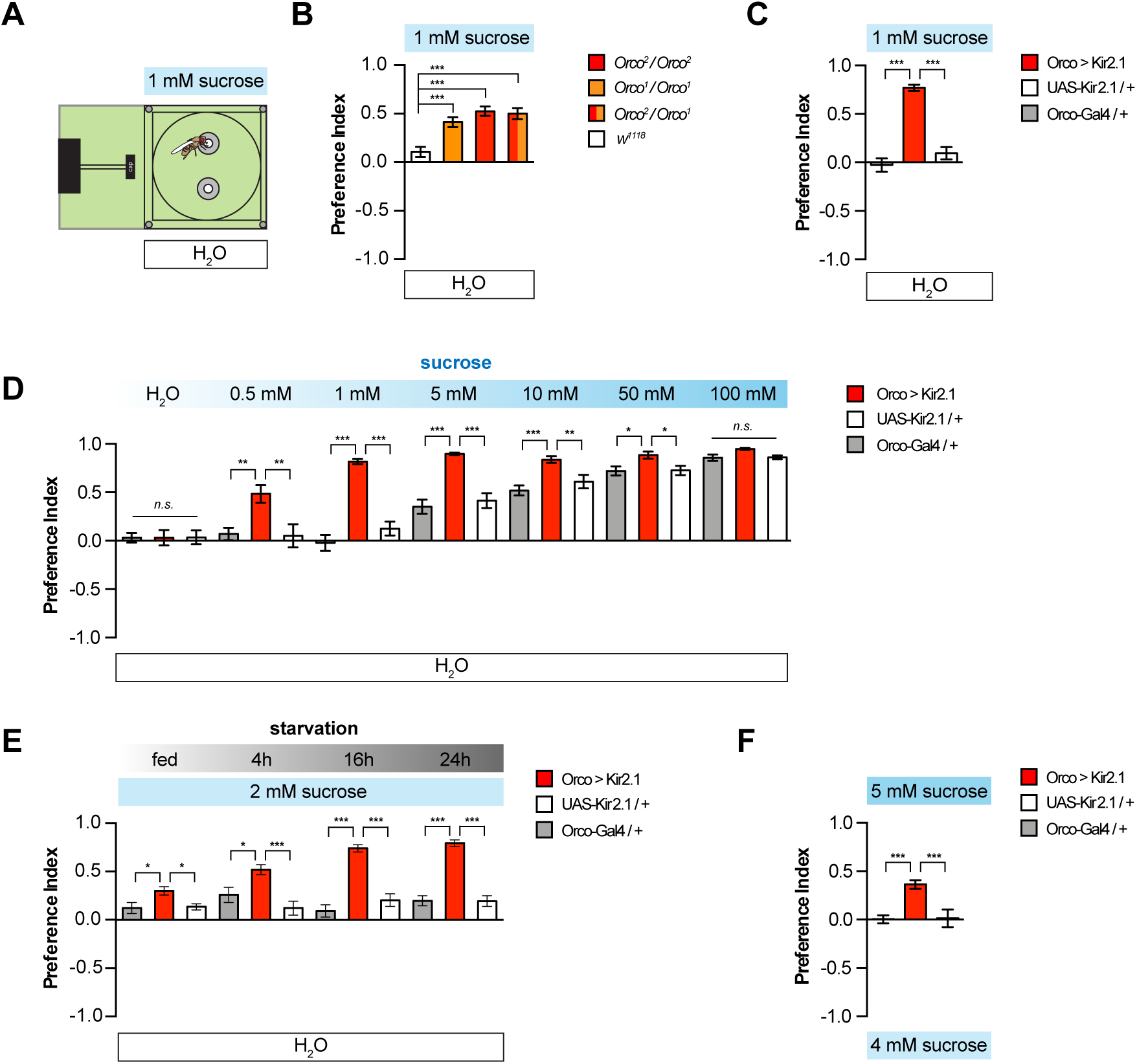
Olfactory impairment enhances sucrose preference and discrimination. (A) Experimental setup of the FlyPAD, where flies were given the choice between 1 mM sucrose and H_2_O, both in 1% agar. (B) Impact of *Orco* mutations on preference between 1 mM sucrose (up) and H_2_O (down), (control, *w*^1118^ (white), *Orco*^1^/*Orco*^1^ (orange), *Orco*^2^/*Orco*^2^ (red), *Orco*^2^/*Orco*^1^ (red/orange) n=21-29. (C) Impact of silencing Orco ORNs with Kir2.1 on preference for 1 mM sucrose over water, n=18-22. (D) Preference of *Orco > Kir2.1* and control flies for increasing concentrations of sucrose (H_2_O, 0.5 mM, 1 mM, 5 mM, 10 mM, 50 mM, 100 mM) and H_2_O n=13-27. (E) Preference of *Orco > Kir2.1* and control flies for 2mM sucrose versus H_2_O under different starvation conditions (fed, 4h, 16h, 24h) n=21-26. (F) Discrimination between 5 mM (positive) and 4 mM (negative) sucrose by *Orco > UAS-Kir2.1* and control flies n=20-26. For all panels, values represent mean +/− SEM; n.s non-significant, * p<0.05, ** p<0.01, *** p<0.001 with 1-way ANOVA and Dunett post hoc test (B) or Tukey HSD post hoc test (C-F).

Internal state strongly impacts taste and feeding, and olfactory loss has been linked to metabolic changes^17^. Therefore, we next asked whether the enhanced sugar feeding of hyposmic flies may be due to increased hunger. We measured preference for 2 mM sucrose over water after varying starvation times, and observed that starvation increased the sugar preference of hyposmic flies but had no impact on the feeding of controls, which were never strongly attracted to this low sugar concentration (Figure 1E,S1D). Moreover, hyposmic flies had significantly higher sucrose preference than controls at all levels of starvation tested from 0 to 24 hours. These results demonstrate that starvation is insufficient to account for the effects of hyposmia and suggest that hyposmic flies are better able to detect low concentrations of sucrose and discriminate them from water. To further explore this idea, we asked whether hyposmia affects the ability of flies to discriminate between very similar sugar concentrations. Indeed, while control flies fed equally on 4 mM and 5 mM sucrose when given the choice between the two, hyposmic flies dramatically preferred the higher concentration (Figure 1F,S1E). Thus, hyposmia not only lowers the threshold for sugar detection, but also enhances sweet taste discrimination.

Although our feeding experiments point towards an effect of hyposmia on taste sensitivity, FlyPAD and other feeding assays introduce the possibility that behavior is influenced by post-ingestive effects. Therefore, we directly measured sweet taste sensitivity using the proboscis extension reflex (PER). Flies detect taste input through different external sensory organs, primarily the tarsi and the labella^18^. Stimulation of either organ with sucrose evokes PER with concentration-dependent probability, and we found that flies with hyposmia from silencing of Orco ORNs exhibited strongly elevated rates of PER at low sucrose concentrations (Figure 2A,B). This effect was independent of starvation, as both fed and starved hyposmic flies displayed increased sucrose sensitivity (Figure 2A-C). Moreover, flies rendered anosmic through the surgical removal of both antenna and maxillary palps also displayed increased PER to sucrose, demonstrating that heightened sweet taste sensitivity is a general consequence of olfactory impairment (Figure 2D). Notably, although it was not the focus, increased sucrose PER in flies with surgically-removed olfactory organs was also incidentally observed in a prior study^19^ To ask whether increasing olfactory input is sufficient to suppress taste sensitivity, we optogenetically activated Orco ORNs and measured the effects on PER^20^. Flies expressing CsChrimson and fed the obligate cofactor all-*trans*-retinal showed lower PER levels in the presence of red light than genetically identical controls which were not fed retinal, demonstrating that manipulating the level of olfactory input can modulate sucrose sensitivity in either direction (Figure 2E). Interestingly, previous work indicates that activating all Orco ORNs carries a positive valence^21^. Thus, we expect that our results are not from a generalized effect on hedonics or a biased cognitive state^22^. Instead, they may highlight a trade-off between taste and smell resulting in a type of homeostatic control over the chemical senses.

**Figure 2.**
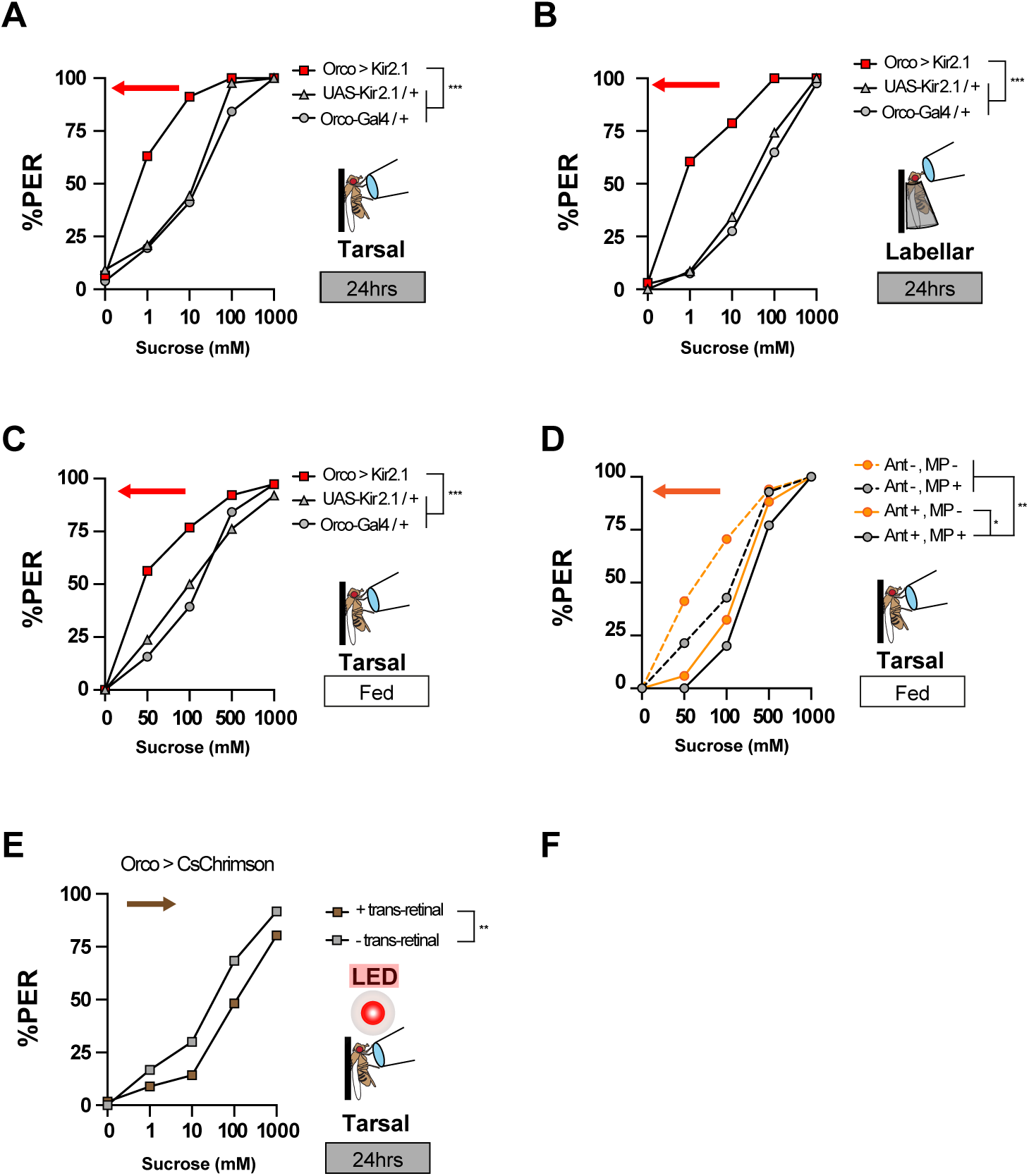
Olfactory impairment increases sucrose responsiveness in PER. (A-C) PER of Orco-silenced and control flies following tarsal (A and C) or labellar (B) stimulation with increasing concentrations of sucrose, in starved (A and B) or fed (C) conditions, n=35-53. (D) Tarsal PER in fed flies following surgical removal of antenna (Ant-, dashed line), maxillary palps (MP-, orange), or both (Ant-, MP-, dashed orange) n=34-35. (E) PER following ORN activation in *Orco > CsChrimson* flies fed all trans retinal (brown squares) compared to controls of the same genotype not fed all trans retinal (grey squares) n=27-36. Arrows point in the direction of shift in sucrose response. Values are total percentage of flies that displayed PER, *p<0.05, **p<0.01, ***p<0.001; 2-way repeated measurements ANOVA with Tukey HSD post hoc test.

If, as we suggest, hyposmia results in increased sweet sensitivity rather than simply elevating feeding responses, we may expect to observe increased sensitivity to other taste modalities, including those that carry a negative valence. We first tested this idea in a traditional bitter sensitivity assay where flies chose between feeding on 4 mM sucrose or 10 mM sucrose mixed with concentrations of bitter L-canavanine (Figure 3A,S2A)^23–25^. Here, ORN-silenced flies displayed dramatically enhanced avoidance of L-canavanine, and this effect was independent of starvation and replicated in *Orco* mutants (Figure 3A and S2B,C). However, formally, this could have resulted from elevated attraction to the plain (4 mM) sucrose when L-canavanine masked the sweetness of the mixture. Therefore, we next tested preference between low concentrations of bitter denatonium and water (Figure 3B,S2D). Once again, hyposmic flies avoided bitter more strongly than controls, suggesting increased bitter sensitivity.

**Figure 3.**
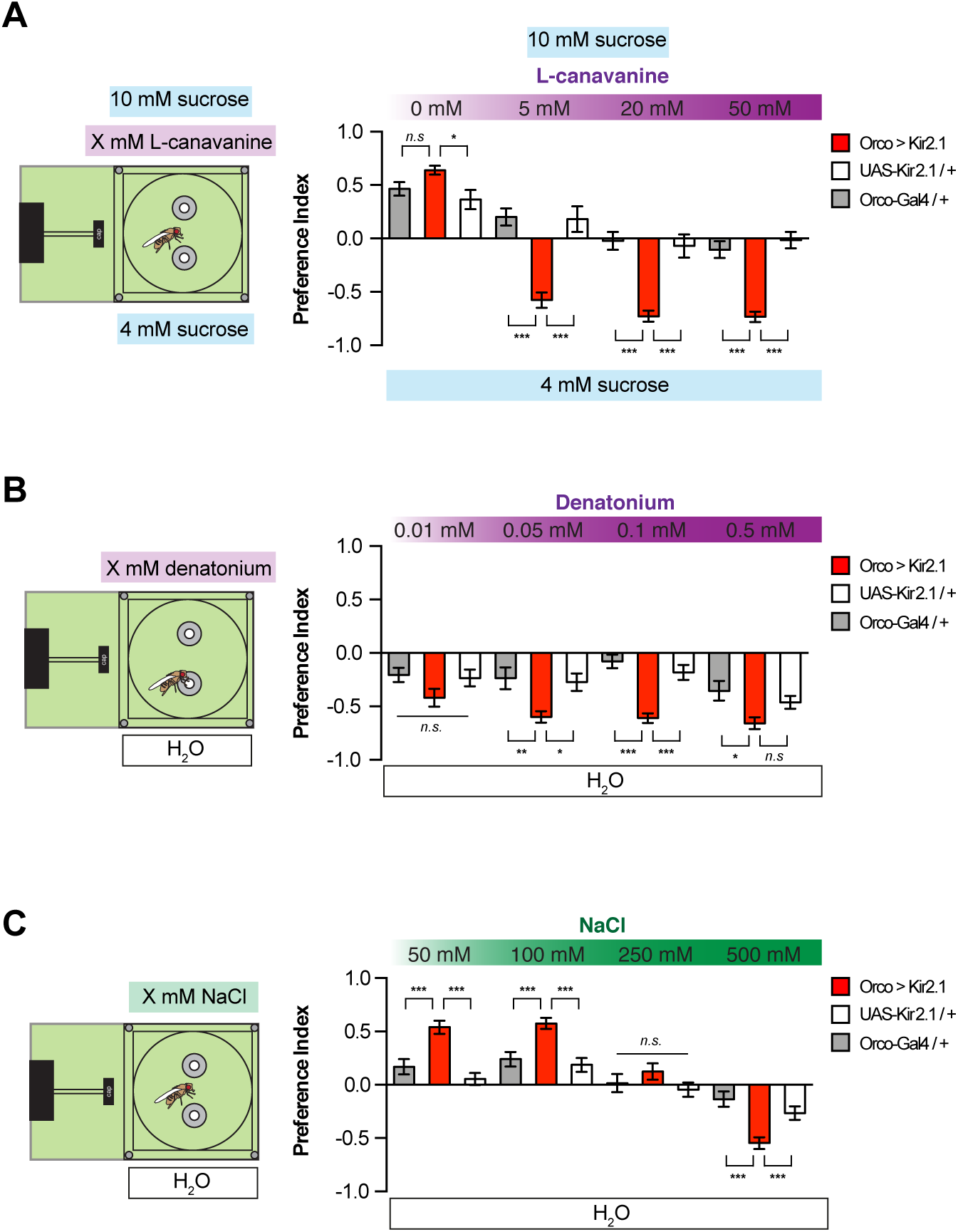
Hyposmia increases responsiveness to bitter and salt. (A-C) Preferences of <u>*Orco > Kir2.1*</u> and control flies in: (A) a choice between increasing concentrations of L-canavanine (0 mM, 5 mM, 20 mM, 50 mM) mixed with 10 mM of sucrose versus 4 mM of sucrose, n=11-29; (B) a choice between increasing concentrations of Denatonium (0.01 mM, 0.05 mM, 0.1 mM, 50 mM) versus water, n=17-25; (C) a choice between increasing concentrations of NaCl (50 mM, 100 mM, 250 mM, 500 mM) versus water, n=19-26. Values are mean +/− SEM, n.s non-significant, * p<0.05, ** p<0.01, *** p<0.001 by 1-way ANOVA with Tukey HSD post hoc test.

Finally, we tested salt preference, which presents an interesting case because NaCl is attractive at low concentrations and aversive at high concentrations^26–28^. Remarkably, hyposmic flies displayed enhanced preference for 50 mM and 100 mM NaCl, and enhanced avoidance of 500 mM NaCl (Figure 3C,S2E). These data fit with the model that olfactory loss affects taste sensitivity across modalities, rather than influencing a particular behavioral output.

In theory, manipulating ORN firing rates could impact taste in a number of different ways. ORNs and olfactory local neurons (LNs) are known to release modulatory neuropeptides, which could impact gustatory receptor neurons (GRNs) or downstream taste circuits from a distance^29,30^. We used calcium imaging to measure sweet GRN activation by sucrose, but saw no effect in *Orco* mutants (Figure 4A,B). Therefore, we suspected that olfactory and gustatory circuits are likely to interact at higher levels in the brain. To explore this possibility, we tested the effect of manipulating olfactory projection neurons (PNs). Silencing of a PN subset labeled by *GH146-Gal4* produced sweet and bitter sensitization that was comparable to ORN silencing (Figure 4C-E,S3A-D). Moreover, optogenetic activation of PNs dramatically suppressed sugar-evoked PER (Figure 4F). Therefore, modulation of taste circuits appears to occur downstream of olfactory PNs. One candidate population for higher-order integration is the PAM dopaminergic neurons of the mushroom body, which respond to sweet stimuli and odours, and are involved in reward coding during appetitive learning^12,13,31–34^. Labellar stimulation with 10 mM sucrose evoked enhanced PAM γ5 activation in ORN-silenced flies compared to controls (Figure 4G,H). However, this difference was eliminated in responses to 1M sucrose, with an opposite trend toward lower activity in the hyposmic flies. Notably, control flies had increased PAM responses at 1M compared to 10 mM, while hyposmic flies showed similar activity across both concentrations. We also observed a higher baseline GCaMP fluorescence of PAM γ5 in the hyposmic individuals, especially in the dorso-medial area (Figure S3E,F). Overall, this could reflect a loss of gain control in the taste system following suppression of olfactory input.

**Figure 4.**
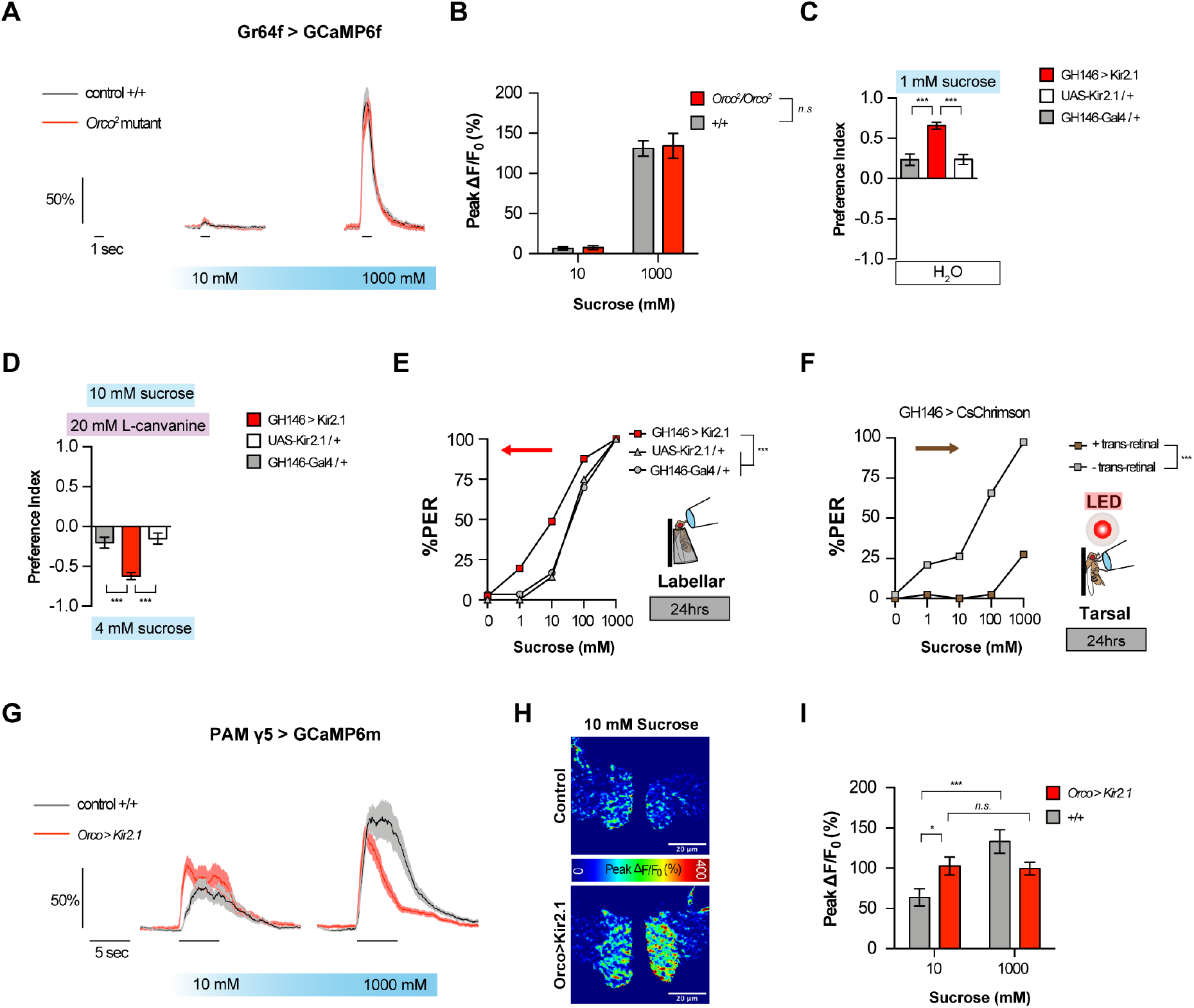
Taste perception enhancement coded in higher order brain areas. (A-B) Calcium imaging of *Gr64f-Gal4* sweet gustatory receptor neurons in response to labellar stimulation with increasing concentrations of sucrose in an *Orco*^2^ homozygote background (red), and controls (grey), showing time curves (A) and peak values (B). Values are mean +/− SEM, n.s.= non-significant; two way repeated measurements ANOVA n=15 (C-D) Preferences of *GH146 > Kir2.1* and control flies in: (C) a choice between 1 mM sucrose versus H_2_O n=33-39; (D) a choice between 20 mM L-canavanine mixed with 10mM of sucrose versus 4mM of sucrose, n=25-31. Values represent mean +/− SEM, ***p<0.001 by 1-way ANOVA, Tukey HSD post hoc test (E) Labellar PER of *GH146 > Kir2.1* and control flies following stimulation with increasing concentrations of sucrose following 24h starvation n=28-41. (F) Tarsal PER following PN activation in *Orco > CsChrimson* flies fed all trans retinal (brown squares) compared to controls of the same genotype not fed all trans retinal (grey squares) n=38-40 Arrows point the shift of sucrose response. Values are total percentage of flies that displayed PER. (G-I) Calcium imaging of PAM γ5 neurons in *Orco > Kir2.1* (red) and wild type background (grey), showing time curves (G), heatmap 10mM (H) and peak values (I) n=62-64. ***p<0.001 with 2-way repeated measures ANOVA with Tukey HSD post hoc test.

Although we do not fully understand the mechanistic connection between olfactory and gustatory circuits that underlies the phenotypes we observed, we can speculate on some possibilities. Second-order olfactory and gustatory neurons both project to the lateral protocerebrum, where cross-talk could easily occur^35^. Even lateral inhibition may be sufficient to mediate suppression of gustatory responses by olfactory activity. The transfer function between ORNs and PNs in the antennal lobe is known to scale inversely with total ORN activity^15,36,37^, and one could imagine a similar gain control mechanism operating between the two chemosensory modalities. This could provide a basic mechanism for attention, where taste becomes more salient in the absence of other food cues to guide feeding decisions. It would be interesting to examine whether cross-modal chemosensory inhibition is bidirectional, by measuring olfactory responses in hypogeusic flies.

Since no odors were presented in any of our experiments, the effects of *Orco* mutations and ORN-silencing on gustatory sensitivity could have arisen from either acute reduction in spontaneous activity within olfactory circuits or chronic olfactory deprivation. Nevertheless, our observation that short-term optogenetic activation of ORNs is sufficient to inhibit PER suggests that olfactory input can acutely suppress sweet taste. However, it is also important to note that not all olfactory activity exerts net suppression on taste sensitivity, as specific odors can either enhance or suppress PER and feeding^38–41^. Thus, complex connections likely exist between olfactory and gustatory circuits that go beyond the scope of what was revealed through our broad manipulations of ORN activity.

## SUPPLEMENTAL INFORMATION

Three supplemental figures can be found in supplemental materials.

## ACKNOWLEDGMENTS

We thank Jinfang Li for pilot data establishing the conditions for imaging and analysis of the PAM. We also thank the Bloomington Stock Center for fly stocks. This work was funded primarily by the Canadian Institutes of Health Research (CIHR) operating grant FDN-148424. M.D.G. is a Michael Smith Foundation for Health Research Scholar.

## AUTHOR CONTRIBUTIONS

PJ conceived of the project, performed all the behavioral experiments, and wrote the first draft of the paper. MS performed the imaging experiments. P-YM participated in experimental design and analysis. MDG supervised the project and co-wrote the paper with PJ.

## DECELARATION OF INTERESTS

The authors declare no conflicts of interest.

## KEY RESOURCES TABLE

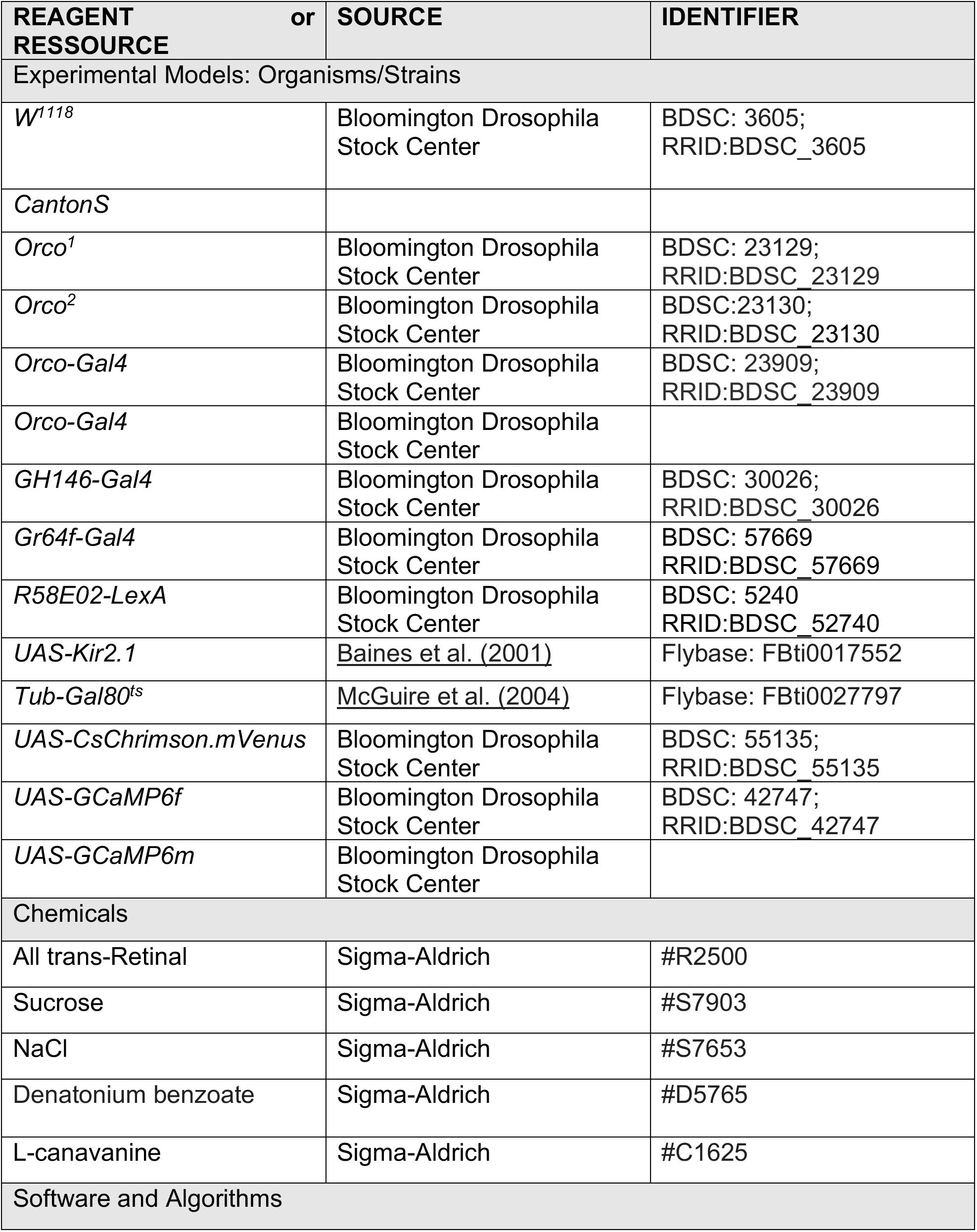
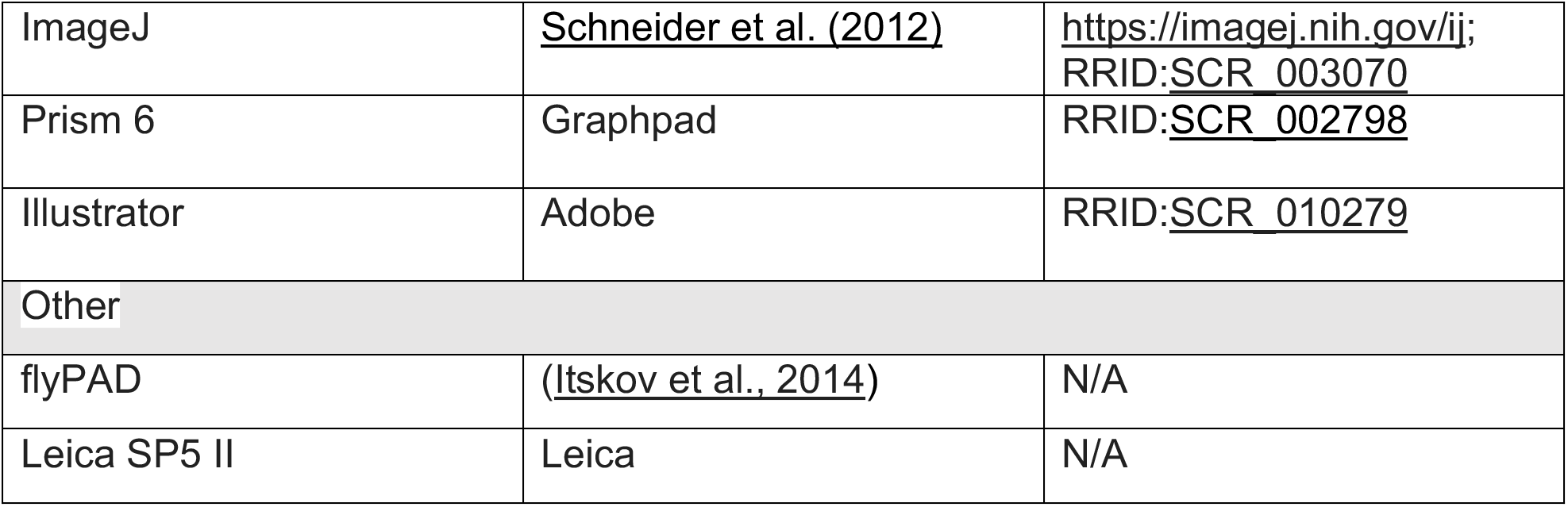

## EXPERIMENTAL MODEL AND SUBJECT DETAILS

The *Drosophila melanogaster* model was used for experimentation, with mutants and transgenic lines detailed in the Key Resources Table.

## METHOD DETAILS

### Fly strains

Fly stocks were reared on standard food at 25°C and 70% humidity under 12/12 hrs light/dark cycle. Genotypes used were: *Orco-GAL4* (BDSC 23909), *GH146-GAL4* (BDSC 30026), *Orco*^2^ (BDSC 23130), *Orco*^1^ (BDSC 23129), *UAS-Kir2.1* (FBti0017552), *Tub-Gal80*^*ts*^ (FBti0027797), *20XUAS-IVS-CSChrimson.mVenus* (BDSC 55135), *UAS-GCaMP6f* (BDSC 42747), and *Gr64f-Gal4* (BDSC 57669), *GH146-Gal4* (BDSC 30026), *R58E02-LexA* (BDSC 5240). All experiments were performed with female flies to reduce variability, given that sex differences were not a subject of investigation.

### FlyPAD experiments

FlyPAD assays were performed similarly to those previously described^16^. Flies were individually transferred to flyPAD arenas by mouth aspiration and allowed to feed for one hour at 25°C, 70% RH. FlyPAD data were acquired using the Bonsai framework^43^, and analyzed in MATLAB using custom-written software^42^. Values for n shown in the figures indicate the number of flies tested. Depending on the experiment, the two channels were filled with different solutions mixed with 1% agar. Details of these mixtures are presented in figures and figure legends. For starvation, flies were kept in vials with 1% agar for the specified amount of time (fed, 8h, 16h, 24h).

### Proboscis extension response assays (PER)

Tarsal PER assays were performed as described previously^44^. 2 – 5 day-old mated females of the indicated genotypes were collected, gently anaesthetized with CO_2_, and fixed by the dorsal thorax onto a glass slide using myristic acid in groups of 20 flies. For labellar PER, flies were mounted inside pipette tips that were cut to size so that only the head was exposed. The tubes were sealed at the end with tape, and positioned on a glass slide with double-sided tape. In both conditions, flies were allowed to recover for 2 hours at room temperature in a humidified-dark box, and stimulations were performed under a dissection microscope. Flies were water satiated by allowing drinking until they no longer responded to water. Stimuli were then presented in series at increasing concentrations using a 20mL pipette attached to a 1 mL syringe. Starved flies were food-deprived in 1% agar vials for the specified amount of time before testing. For photoactivation of ORNs and PNs, the stimulations were applied under a red LED.

For olfactory organ removal, the third antennal segments and/or maxillary palps were removed with forceps while flies were anesthetized on a CO_2_ pad. Surgery was performed prior to starvation, and flies were given ~30 min to recover in food vials before starvation. Control flies with intact olfactory organs were anesthetized for the same duration as antennectomized flies. In this experiment, *CantonS* flies were used because *w*^1118^ flies did not recover well from surgery.

### Calcium imaging

For calcium imaging of sweet GRNs, 2 – 5 day-old female flies were briefly anesthetized using CO_2_, and placed in custom chamber suspended from their cervix. To ensure immobilization, a small drop of nail polish was applied to the back of the neck and the proboscis was pulled to extension and waxed out on both sides. A modified dental waxer was used to apply wax on each side of the chamber rim, making little contact with the feeding structure. Flies were left to recover in a humidified chamber for 1 hr. A small window of cuticle was removed from the top of the head, exposing the SEZ. Adult Hemolymph Like (AHL) buffer was immediately applied to the preparation (108 mM NaCl, 5 mM KCl, 4 mM NaHCO3, 1 mM NaH2PO4, 5 mM HEPES, 15 mM ribose, pH 7.5). The air sacs, fat, and esophagus were clipped and removed to allow clear visualization on the SEZ. Once ready to image, AHL buffer was added that includes Mg^2+^ and Ca^2+^ (108 mM NaCl, 5 mM KCl, 4 mM NaHCO3, 1 mM NaH2PO4, 5 mM HEPES, 15 mM ribose, 2 mM Ca^2+^, and 8.2 mM Mg^2+^). GCaMP6f fluorescence was observed using a Leica SP5 II Confocal microscope with a 25× water immersion objective. The relevant area of the SEZ was visualized at a zoom of 4×, a line speed of 8000 Hz, a line accumulation of 2, and resolution of 512 × 512 pixels. The pinhole was opened to 2.98 AU. For each taste stimulation, data was acquired during a baseline of 5 s prior to stimulation, 1 s during tastant application, and 9 s following the stimulation. Tastant stimulations were done using a pulled capillary pipette that was filed down to match the size of the proboscis and fit over all taste sensilla on both labellar palps. The pipette was filled with 1 – 2 μl of a tastant and positioned close to the proboscis labellum. At 5 s a micromanipulator was used to apply the tastant to the labellum manually. Between taste stimulations of differing solutions, the pipette was washed with water. Sucrose solutions were applied in the order of increasing concentration.The maximum change in fluorescence (ΔF/F) was calculated using the peak intensity (average of 3 time points) minus the average intensity at baseline (average of 10 time points), divided by the baseline.

*In vivo* calcium imaging of gamma lobe PAM neurons was performed the same way as for GRNs but with the following differences: additional cuticle was removed for full visualization of the PAM brain region and there was no need to cut the esophagus. In addition, in the LAS AF program, line accumulation was set to 1 (as opposed to 2 for GRN imaging), the pinhole was opened to 80 micrometers, and the gain remained constant for each fly. Each PAM recording consisted of 10 s of baseline, 5 s of labellar stimulation, and 15 s after the stimulus was removed. Based on pilot imaging, we established conditions for excluding flies that did not respond robustly to stimulation. Flies were excluded if they did not show a clear visual response to 10 mM sucrose *and* the response to 1 M was less than 50%. In a few instances, large, spontaneous oscillations or significant brain movement made it impossible to determine the change in fluorescence due to labellar stimulation, so these flies were also removed from the final dataset. Left and right gamma lobes were analyzed and plotted individually^34^, so the final N represents regions of interest (ROIs), for which there were two per fly, similar to a previous report^34^. While performing these experiments, we noted subtle differences in the baseline fluorescence between genotypes. In particular, the *Orco>Kir2.1* flies had pronounced baseline fluorescence in the superior region of the gamma lobe, so we quantified baseline fluorescence from the same 10 points in the full ROI, an ROI that excluded this superior region, and an ROI that only included this superior region. We compared peak ΔF/F_0_ sucrose responses in each ROI for a subset of flies, but this did not change the data trends in any significant way, so we continued only analyzing ΔF/F_0_ sucrose responses in the full ROI.

## QUANTIFICATION AND STATISTICAL ANALYSIS

Statistical tests were performed using GraphPad Prism 6 software. Descriptions and results of each test are provided in the figure legends. Sample sizes are indicated in the figure legends. Sample sizes were determined prior to experimentation based on the variance and effect sizes seen in prior experiments of similar types. All replicates were biological replicates using different individual flies. Data for all quantitative experiments were collected on at least three different days, and behavioral experiments were performed with flies from at least two independent crosses. Specific definitions of replicates are as follows. For PER, each replicate is composed of 20 independent flies tested in parallel. PER response was calculated as a percentage of proboscis extensions following tastant stimulation of the tarsi or the labellum. For flyPAD experiments, each data point is the calculated preference of an individual fly over the course of the experiment. Preference index (PI) was calculated as ((number of sip Left side) – (number of sip Right side))/(total number of sips). For flyPAD experiments, the data from individual flies were removed if the fly did not pass a set minimum threshold of sips (15), or the data showed hallmarks of a technical malfunction (rare). All the quantitative data used for statistical tests can be found as supplements for each figure. Repeated measurement ANOVA were performed on PER data, with Tukey HSD posthoc test^45^. For flyPAD analyses, one way ANOVA with a Dunett posthoc was used for the *Orco* mutant experiments and a Tukey posthoc was used for the others in which the group of interests were compared to their two parental controls. Two-way ANOVA and Bonferroni posthoc test were used on the imaging data.

## Notes

### Competing Interest Statement

The authors have declared no competing interest.

## REFERENCES

1. Galton, F. INQUIRIES INTO HUMAN FACULTY. 305.

2. Zopf, G.W. (1963). Sensory Homeostasis. In Progress in Brain Research (Elsevier), pp. 114–121.

3. Bavelier, D., and Neville, H.J. (2002). Cross-modal plasticity: where and how? Nat Rev Neurosci 3, 443–452.

4. Kupers, R., and Ptito, M. (2014). Compensatory plasticity and cross-modal reorganization following early visual deprivation. Neuroscience & Biobehavioral Reviews 41, 36–52.

5. Blankenship, M.L., Grigorova, M., Katz, D.B., and Maier, J.X. (2019). Retronasal Odor Perception Requires Taste Cortex, but Orthonasal Does Not. Current Biology 29, 62–69.e3.

6. Shimojo, S. (2001). Sensory modalities are not separate modalities: plasticity and interactions. Current Opinion in Neurobiology 11, 505–509.

7. Landis, B.N., Scheibe, M., Weber, C., Berger, R., Brämerson, A., Bende, M., Nordin, S., and Hummel, T. (2010). Chemosensory interaction: acquired olfactory impairment is associated with decreased taste function. J Neurol 257, 1303–1308.

8. Reichert, J.L., and Schöpf, V. (2018). Olfactory Loss and Regain: Lessons for Neuroplasticity. Neuroscientist 24, 22–35.

9. Frasnelli, J., Collignon, O., Voss, P., and Lepore, F. (2011). Crossmodal plasticity in sensory loss. In Progress in Brain Research (Elsevier), pp. 233–249.

10. Vosshall, L.B., and Stocker, R.F. (2007). Molecular Architecture of Smell and Taste in *Drosophila*. Annu. Rev. Neurosci. 30, 505–533.

11. Scott, K. (2018). Gustatory Processing in *Drosophila melanogaster*. Annu. Rev. Entomol. 63, 15–30.

12. Yamagata, N., Ichinose, T., Aso, Y., Plaçais, P.-Y., Friedrich, A.B., Sima, R.J., Preat, T., Rubin, G.M., and Tanimoto, H. (2015). Distinct dopamine neurons mediate reward signals for short- and long-term memories. Proc Natl Acad Sci USA 112, 578–583.

13. Shyu, W.-H., Chiu, T.-H., Chiang, M.-H., Cheng, Y.-C., Tsai, Y.-L., Fu, T.-F., Wu, T., and Wu, C.-L. (2017). Neural circuits for long-term water-reward memory processing in thirsty Drosophila. Nat Commun 8, 15230.

14. Larsson, M.C., Domingos, A.I., Jones, W.D., Chiappe, M.E., Amrein, H., and Vosshall, L.B. (2004). Or83b Encodes a Broadly Expressed Odorant Receptor Essential for Drosophila Olfaction. Neuron 43, 703–714.

15. Cao, L.-H., Yang, D., Wu, W., Zeng, X., Jing, B.-Y., Li, M.-T., Qin, S., Tang, C., Tu, Y., and Luo, D.-G. (2017). Odor-evoked inhibition of olfactory sensory neurons drives olfactory perception in Drosophila. Nat Commun 8, 1357.

16. Itskov, P.M. (2014). Automated monitoring and quantitative analysis of feeding behaviour in Drosophila. NATURE COMMUNICATIONS, 10.

17. Libert, S., Zwiener, J., Chu, X., VanVoorhies, W., Roman, G., and Pletcher, S.D. (2007). Regulation of Drosophila Life Span by Olfaction and Food-Derived Odors. Science 315, 1133–1137.

18. Thoma, V., Knapek, S., Arai, S., Hartl, M., Kohsaka, H., Sirigrivatanawong, P., Abe, A., Hashimoto, K., and Tanimoto, H. (2016). Functional dissociation in sweet taste receptor neurons between and within taste organs of Drosophila. Nat Commun 7, 10678.

19. Devineni, A.V., Sun, B., Zhukovskaya, A., and Axel, R. (2019). Acetic acid activates distinct taste pathways in Drosophila to elicit opposing, state-dependent feeding responses. eLife 8, e47677.

20. Klapoetke, N.C., Murata, Y., Kim, S.S., Pulver, S.R., Birdsey-Benson, A., Cho, Y.K., Morimoto, T.K., Chuong, A.S., Carpenter, E.J., Tian, Z., et al. (2014). Independent optical excitation of distinct neural populations. Nat Methods 11, 338–346.

21. Tao, L., Ozarkar, S., and Bhandawat, V. (2020). Mechanisms underlying attraction to odors in walking Drosophila. PLoS Comput Biol 16, e1007718.

22. Bateson, M., Desire, S., Gartside, S.E., and Wright, G.A. (2011). Agitated Honeybees Exhibit Pessimistic Cognitive Biases. Current Biology 21, 1070–1073.

23. Shim, J., Lee, Y., Jeong, Y.T., Kim, Y., Lee, M.G., Montell, C., and Moon, S.J. (2015). The full repertoire of Drosophila gustatory receptors for detecting an aversive compound. Nat Commun 6, 8867.

24. Lee, Y., Kang, M.J., Shim, J., Cheong, C.U., Moon, S.J., and Montell, C. (2012). Gustatory Receptors Required for Avoiding the Insecticide L-Canavanine. Journal of Neuroscience 32, 1429–1435.

25. Mitri, C., Soustelle, L., Framery, B., Bockaert, J., Parmentier, M.-L., and Grau, Y. (2009). Plant Insecticide L-Canavanine Repels Drosophila via the Insect Orphan GPCR DmX. PLoS Biol 7, e1000147.

26. Zhang, Y.V., Ni, J., and Montell, C. (2013). The Molecular Basis for Attractive Salt-Taste Coding in Drosophila. Science 340, 1334–1338.

27. Jaeger, A.H., Stanley, M., Weiss, Z.F., Musso, P.-Y., Chan, R.C., Zhang, H., Feldman-Kiss, D., and Gordon, M.D. (2018). A complex peripheral code for salt taste in Drosophila. eLife 7, e37167.

28. Chen, Y.-C.D., Park, S.J., Joseph, R.M., Ja, W.W., and Dahanukar, A.A. (2019). Combinatorial Pharyngeal Taste Coding for Feeding Avoidance in Adult Drosophila. Cell Reports 29, 961–973.e4.

29. Carlsson, M.A., Diesner, M., Schachtner, J., and Nässel, D.R. (2010). Multiple neuropeptides in the Drosophila antennal lobe suggest complex modulatory circuits. J. Comp. Neurol. 518, 3359–3380.

30. Nässel, D.R., and Zandawala, M. (2019). Recent advances in neuropeptide signaling in Drosophila, from genes to physiology and behavior. Progress in Neurobiology 179, 101607.

31. Yamagata, N., Hiroi, M., Kondo, S., Abe, A., and Tanimoto, H. (2016). Suppression of Dopamine Neurons Mediates Reward. PLoS Biol 14, e1002586.

32. Musso, P.-Y., Lampin-Saint-Amaux, A., Tchenio, P., and Preat, T. (2017). Ingestion of artificial sweeteners leads to caloric frustration memory in Drosophila. Nat Commun 8, 1803.

33. Lewis, L.P.C., Siju, K.P., Aso, Y., Friedrich, A.B., Bulteel, A.J.B., Rubin, G.M., and Grunwald Kadow, I.C. (2015). A Higher Brain Circuit for Immediate Integration of Conflicting Sensory Information in Drosophila. Current Biology 25, 2203–2214.

34. May, C.E., Rosander, J., Gottfried, J., Dennis, E., and Dus, M. (2020). Dietary sugar inhibits satiation by decreasing the central processing of sweet taste. eLife 9, e54530.

35. Talay, M., Richman, E.B., Snell, N.J., Hartmann, G.G., Fisher, J.D., Sorkaç, A., Santoyo, J.F., Chou-Freed, C., Nair, N., Johnson, M., et al. (2017). Transsynaptic Mapping of Second-Order Taste Neurons in Flies by trans-Tango. Neuron 96, 783–795.e4.

36. Olsen, S.R., and Wilson, R.I. (2008). Lateral presynaptic inhibition mediates gain control in an olfactory circuit. Nature 452, 956–960.

37. Wilson, R.I. (2013). Early Olfactory Processing in *Drosophila*: Mechanisms and Principles. Annu. Rev. Neurosci. 36, 217–241.

38. Shiraiwa, T. (2008). Multimodal Chemosensory Integration through the Maxillary Palp in Drosophila. PLoS ONE 3, e2191.

39. Stensmyr, M.C., Dweck, H.K.M., Farhan, A., Ibba, I., Strutz, A., Mukunda, L., Linz, J., Grabe, V., Steck, K., Lavista-Llanos, S., et al. (2012). A Conserved Dedicated Olfactory Circuit for Detecting Harmful Microbes in Drosophila. Cell 151, 1345–1357.

40. Reisenman, C.E., and Scott, K. (2019). Food-derived volatiles enhance consumption in *Drosophila melanogaster*. J Exp Biol 222, jeb202762.

41. Oh, S.M., Jeong, K., Seo, J.T., and Moon, S.J. (2021). Multisensory interactions regulate feeding behavior in *Drosophila*. Proc Natl Acad Sci USA 118, e2004523118.

42. Musso, P.-Y., Junca, P., Jelen, M., Feldman-Kiss, D., Zhang, H., Chan, R.C., and Gordon, M.D. (2019). Closed-loop optogenetic activation of peripheral or central neurons modulates feeding in freely moving Drosophila. eLife 8, e45636.

43. Lopes, G., Bonacchi, N., FrazÃ£o, J., Neto, J.P., Atallah, B.V., Soares, S., Moreira, L., Matias, S., Itskov, P.M., Correia, P.A., et al. (2015). Bonsai: an event-based framework for processing and controlling data streams. Front. Neuroinform. 9.

44. Walker, S.J., Corrales-Carvajal, V.M., and Ribeiro, C. (2015). Postmating Circuitry Modulates Salt Taste Processing to Increase Reproductive Output in Drosophila. Current Biology 25, 2621–2630.

45. Lunney, G.H. (1970). USING ANALYSIS OF VARIANCE WITH A DICHOTOMOUS DEPENDENT VARIABLE: AN EMPIRICAL STUDY1. J Educational Measurement 7, 263–269.

