## Supplemental figures for "Modulation of taste sensitivity by the olfactory system in *Drosophila*"

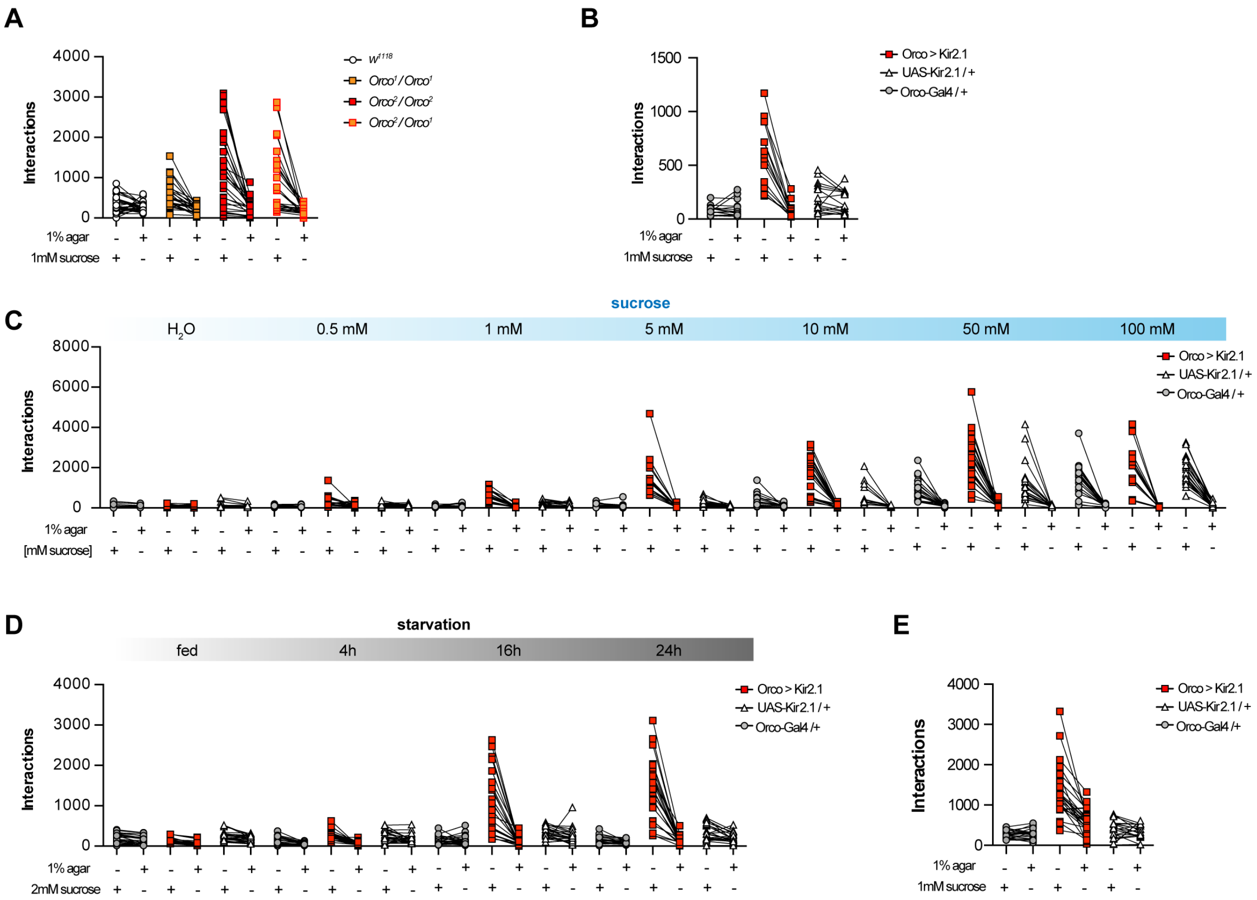


**Figure S1. Olfactory impairment enhances sucrose preference and discrimination. Related to Figure 1.** (A-E) Interaction numbers for each individual fly from experiments shown in Figure 1: Impact of *Orco* mutations (A; Figure 1B) or *Orco-Gal4* silencing (B; Figure 1C) on preference between 1 mM sucrose (left) and $H_2O$ (right), n=21-29; *Orco* > *Kir2.1* flies given choice between increasing concentrations of sucrose ($H_2O$, 0.5 mM, 1 mM, 5 mM, 10 mM, 50 mM, 100 mM) and $H_2O$ (C; Figure 1D), n=13-27; *Orco* > *Kir2.1* flies choosing between 2mM sucrose and $H_2O$ after different starvation times (D; Figure 1E) n=21-26; *Orco* > *Kir2.1* flies choosing between 5 mM and 4 mM sucrose (E; Figure 1F), n=20-26. *Orco* > *Kir2.1* (red squares), *UAS-Kir2.1* / + (white triangles), *Orco-Gal4* / + (grey circles).

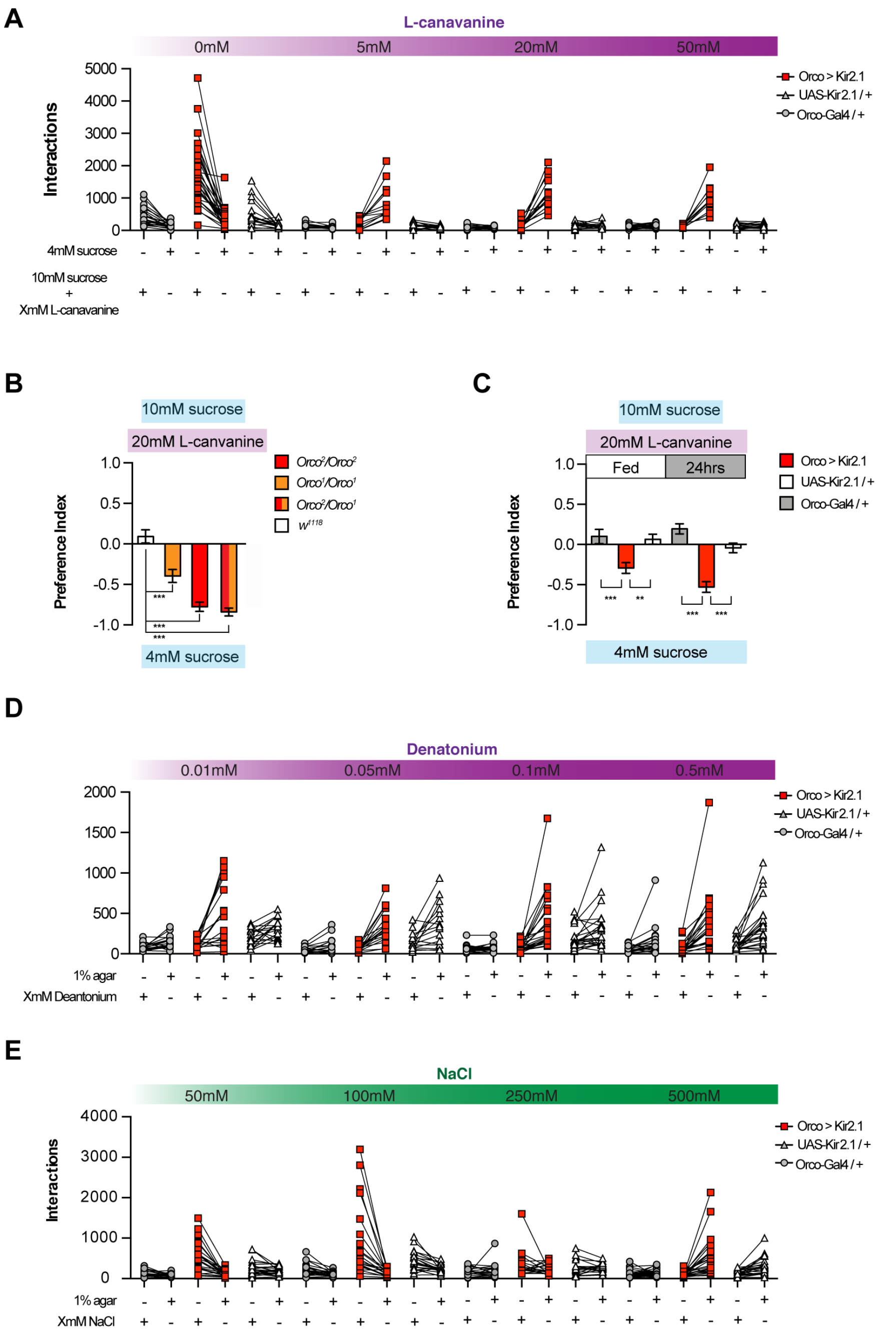


**Figure S2. Loss of smell increases responsiveness to bitter and salt. Related to Figure 3.** (A) Interaction numbers for individual flies in the experiment to measure L-canavanine avoidance after Orco silencing (preferences in Figure 3A), n=11-29. (B) Impact of Orco mutations on preference between 20 mM L-canavanine mixed with 10 mM sucrose versus 4 mM sucrose, (control, $w^{1118}$ (white), *Orco*$^1$/*Orco*$^1$ (orange), *Orco*$^2$/*Orco*$^2$ (red), *Orco*$^2$/*Orco*$^1$ (red/orange) n=[21-23]. (C) Impact of starvation (fed, 24hrs) in the same conditions as (B) *Orco* > *Kir2.1* (red), *UAS-Kir2.1* / + (white), *Orco-Gal4* / + (grey) n=[17-21]. (D and E) Interaction numbers from experiments measuring impact of *Orco-Gal4* silencing on: a choice between increasing concentrations of Denatonium (0.01 mM, 0.05 mM, 0.1 mM, 50 mM) and water (D; preferences shown in Figure 3B), n=17-25; a choice between increasing concentrations of NaCl (50 mM, 100 mM, 250 mM, 500 mM) and water (E; preferences shown in Figure 3C), n=19-26. For (A, D and E) *Orco* > *Kir2.1* (red squares), *UAS-Kir2.1* / + (white triangle), *Orco-Gal4* / + (grey round). *** p<0.001, 1 way ANOVA with Tukey HSD post hoc test (B and C)

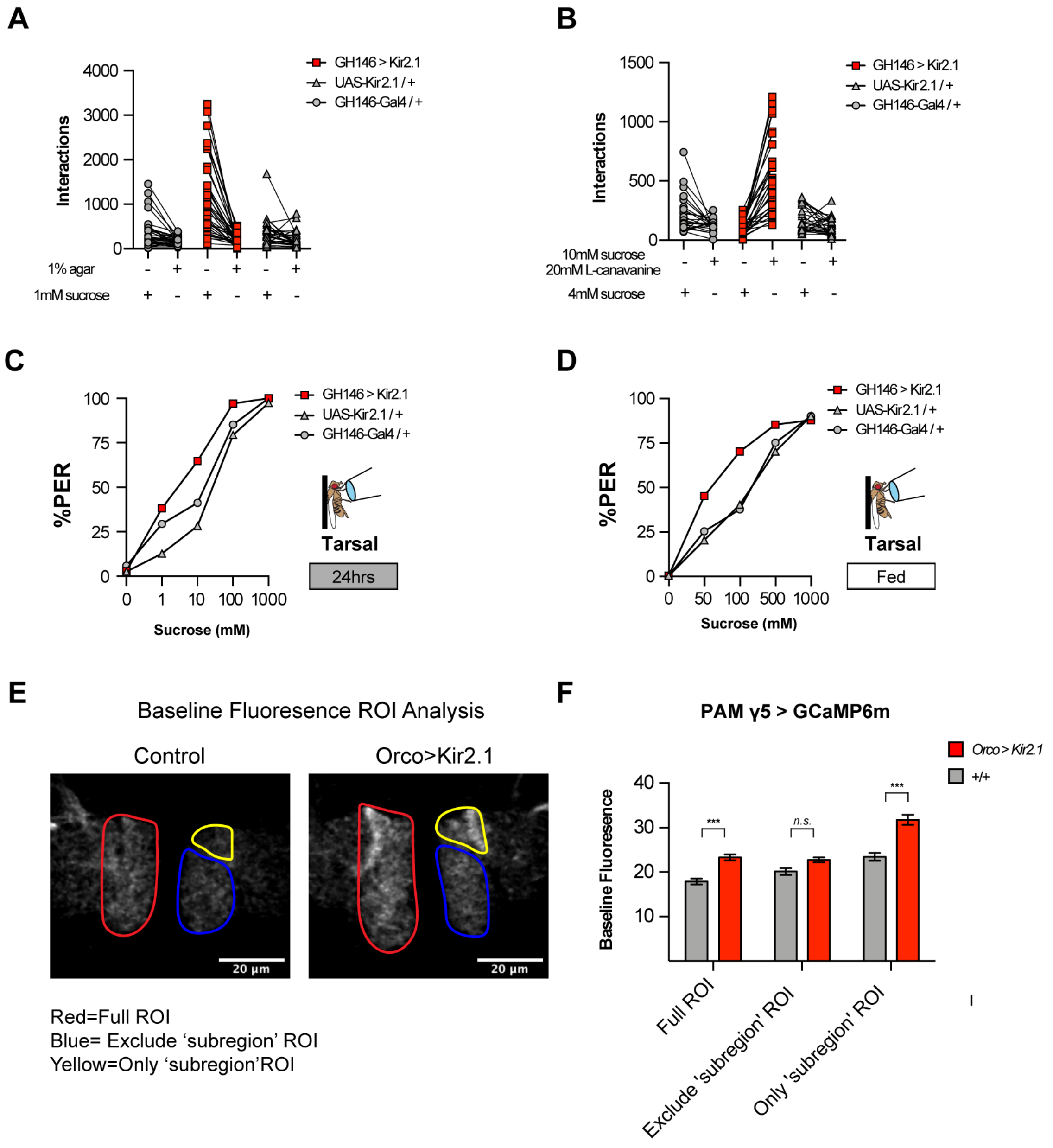


**Figure S3. Impairment of olfactory projection neurons increases taste sensitivity. Related to Figure 4.** (A and B) Interaction numbers for each individual fly from experiments shown in Figure 4: Impact of *GH146-Gal4* silencing on preference between 1 mM sucrose (left) and $H_2O$ (right) (A; Figure 1A), n=33-39; Impact of *GH146-Gal4* silencing on preference between 20 mM L-canavanine mixed with 10 mM sucrose versus 4 mM sucrose (B; Figure 1B), n=25-29. (C and D) Impact of PN silencing on PER following tarsal stimulation of increasing concentrations of sucrose, in 24h starved (C) or fed (D) conditions. *GH146 > Kir2.1* (red squares), *UAS-Kir2.1* / + (white triangles), *GH146-Gal4* / + (grey circles). (E-F) Baseline activity of Different subregion of PAM γ5 neurons in Orco > Kir2.1 and control individuals n=62-64, n.s non-significant, ***p<0.001 by 2 ways ANOVA with Tukey HSD post hoc test (C and D).
